# Gα_i_-derived peptide binds the µ-opioid receptor

**DOI:** 10.1101/729244

**Authors:** Piotr Kosson, Jolanta Dyniewicz, Piotr F. J. Lipiński, Joanna Matalinska, Aleksandra Misicka, Andrzej J. Bojarski, Stefan Mordalski

## Abstract

G protein-coupled receptors (GPCRs) transduce external stimuli into the cell by G proteins via an allosteric mechanism. Agonist binding to the receptor stimulates GDP/GTP exchange within the heterotrimeric G protein complex. Whereas recent structures of GPCR-G protein complexes revealed that the H5, S1 and S2 domains of Gα are involved in binding the active receptor, earlier studies showed that a short peptide analogue derived from the C-terminus (H5) of the G protein transducin (G_t_) is sufficient to stabilize rhodopsin in an active form. Here, we show that a Gα_i_-derived peptide of 12 amino acids binds the µ-opioid receptor (µOR) and acts as an allosteric modulator. The Gα_i_-derived peptide increases µOR affinity for its agonist morphine in a dose-dependent way. These results indicate that the GPCR-Gα peptide interaction observed so far for only rhodopsin can be extrapolated to µOR. In addition, we show that the C-terminal peptide of the Gα_i_ subunit is sufficient to stabilize the active conformation of the receptor. Our approach opens the possibility to investigate the GPCR-G protein interface with peptide modification.

## Introduction

G protein-coupled receptors (GPCRs) are a superfamily of approximately 800 receptors (in humans) transmitting extracellular stimuli (e.g., small molecules, peptides, lipids or light) into the cell. Due to their key regulatory roles in multiple cell types and tissues and their resulting potential for drug targeting, GPCRs have become the most frequently targeted protein class on the market.^1^ On the other hand, the active GPCR signal is transduced into the cell by only 16 different G proteins.^2^ Understanding the molecular basis of GPCR-G protein pairing is essential to understanding the phenomena of functional selectivity and biased signalling, where the type of ligand bound determines which signal transducers are activated by the receptor. Utilizing this knowledge can have significant therapeutic implications, as it may lead to novel drugs with reduced side effects.^3,4^

GPCR activation triggered by extracellular orthosteric agonists through a cascade of conformational changes opens the intracellular side of the receptor for interaction with the heterotrimeric G protein. Experimental evidence supports the model of ternary complex^5^ formation by the receptor, agonist and signal transducer, where the latter stabilizes the active conformation of the receptor.^6^ In this model, which is further supported by NMR studies,^7^ the unbound receptor occupies a certain conformational space including the conformations corresponding to the active state (thus explaining the phenomenon of basal activity). Binding of the agonist to the receptor limits this conformational space but leaves some degree of flexibility that is further narrowed by G protein coupling. Recent functional studies shed light on the process of ternary complex formation. Work by DeVree *et al*.^8^ postulates G protein-dependent low- and high-agonist-affinity states of the receptor. Binding of the agonist promotes the interaction between the receptor and G protein (low-affinity state). Release of GDP by the G protein enables stronger interactions with the GPCR, stabilizing its active state, allosterically affecting the orthosteric (extracellular) region of the receptor, and enhancing the affinity of the agonist (high-affinity state).

Multiple pharmacological and structural studies indicate that other binding partners, including antibodies and nanobodies,^9^ mini-G proteins,^10^ and peptides derived from G_t_^11^ or arrestin-1,^12^ can mimic the effect of G protein binding and thus be used for investigation of GPCR activation. Among these, synthetic peptides derived from the C-terminus of Gα (G-peptides) offer a promising avenue for investigating GPCR activation^13^, and with peptide mutations increasing the affinity to the target receptor^14^, they open a possibility for studying G protein subtype selectivity and identification of the GPCR-G protein interaction hotspots. The crystal structures of active rhodopsin have employed a G protein-derived peptide, and this peptide binds within the intracellular crevice of rhodopsin.^11,15^ Here, our aim was to investigate whether another GPCR in addition to rhodopsin could be bound and stabilized by a peptide derived from a G protein other than G_t_.

## Results

### Target selection

To determine if small peptides derived from other G proteins might be used to study GPCRs other than rhodopsin, we began by identifying the structural and topological features of rhodopsin differentiating it from other crystallized GPCRs. As there are no clear sequence motifs corresponding to matching different G proteins,^16^ we looked at the more general level – i.e., the size of the intracellular loops of the receptors. It appears that intracellular loop 3 (ICL3) of rhodopsin is significantly shorter than those of other GPCRs crystallized in the active state (Figure 1a). Notably, short loops are actually more common than long loops among GPCRs in general, and those with a longer ICL3 are disproportionately over-represented in the small group of GPCRs that have been crystallized (Figure 1b). Given that fact, it should be possible to observe G-peptide binding to a receptor with an ICL3 of similar length to that of rhodopsin. We selected µ-opioid receptor (µOR) for the study. It not only matches the requirement for ICL3 length (it contains 8 residues) but also has a number of well-studied ligands with distinct functional profiles, allowing for more comprehensive experimental studies.

**Figure 1.**
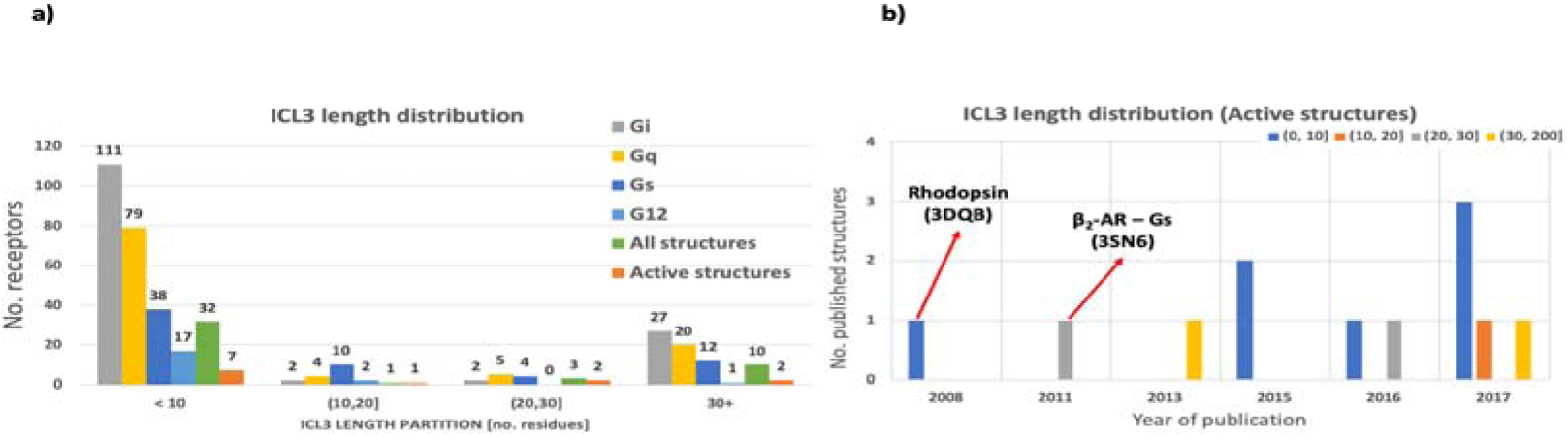
The distribution of the length of ICL3 in GPCRs. **a**, The ICL3 among the GPCRs with known G protein coupling (GPCRdb and IUPHAR GuideToPharmacology data) tends to be short (shorter than 10 amino acids) for the majority of the analysed targets. This regularity is not maintained for G_s_-coupled receptors, with receptor ICL3s spanning all lengths. In the overall pool of GPCRs, the short loops are dominant (ca. 75% of receptors have an ICL3 of ten residues or shorter), whereas the crystallized targets have an overrepresentation of long ICL3s, which might affect the potential binding of a G-peptide. The lengths of ICL3s were calculated based on GPCRdb annotation of the receptor regions. **b**, An analysis of the active-state structures of GPCRs indicates the overrepresentation of the long-ICL3 receptors among the crystalized targets. In addition, rhodopsin, for which the first active-state structure was solved, is also the only target that has been crystalized without an antibody stabilizing the active conformation of the 7TM bundle.

### *In vitro* studies

The binding of the G-peptide was experimentally verified *in vitro*. We used three compounds with different functional profiles: morphine, prototyping an unbiased ligand;^17^ a G_i_-biased agonist PZM21;^3^ and fentanyl, with a strong preference towards β-arrestin recruitment.^18^ Previous studies on G proteins and nanobodies demonstrated the allosteric effect of binding an intracellular agent stabilizing an active conformation,^8,19–21^ resulting in increased agonist affinity for a receptor bound by G protein or nanobodies. Here, we capitalized on the same assumption, observing the changes in the results of a radioligand-binding assay. The results of *in vitro* experiments showed an increase in the affinity of morphine after adding the G-peptide. The leftward shift in the binding curve and the increased amount of radioligand bound indicate the allosteric effect of the added peptide (Figure 2a, 2e). On the other hand, the same experiment with the G protein-biased agonist PZM21 resulted in an increase in the affinity towards µOR but no change in the amount of radioligand bound (Figure 2b, 2f). Finally, the binding affinity of fentanyl was not affected by the G-peptide, but we observed a significantly increased amount of radioligand bound (Figure 2c, 2g). At the same time, the peptide itself did not interact directly with the orthosteric binding site. A binding experiment performed for the G-peptide allowed us to estimate the affinity of the peptide in the micromolar range, which was four orders of magnitude lower than those measured for the reference ligands (Supplementary Figure 1). Saturation experiments performed for [H^3^]DAMGO confirmed the dose dependency of the affinity increase caused by the G-peptide, providing additional confirmation that the allosteric effects we observed in the binding studies are indeed caused by the peptide (Supplementary Figure 2).

**Figure 2.**
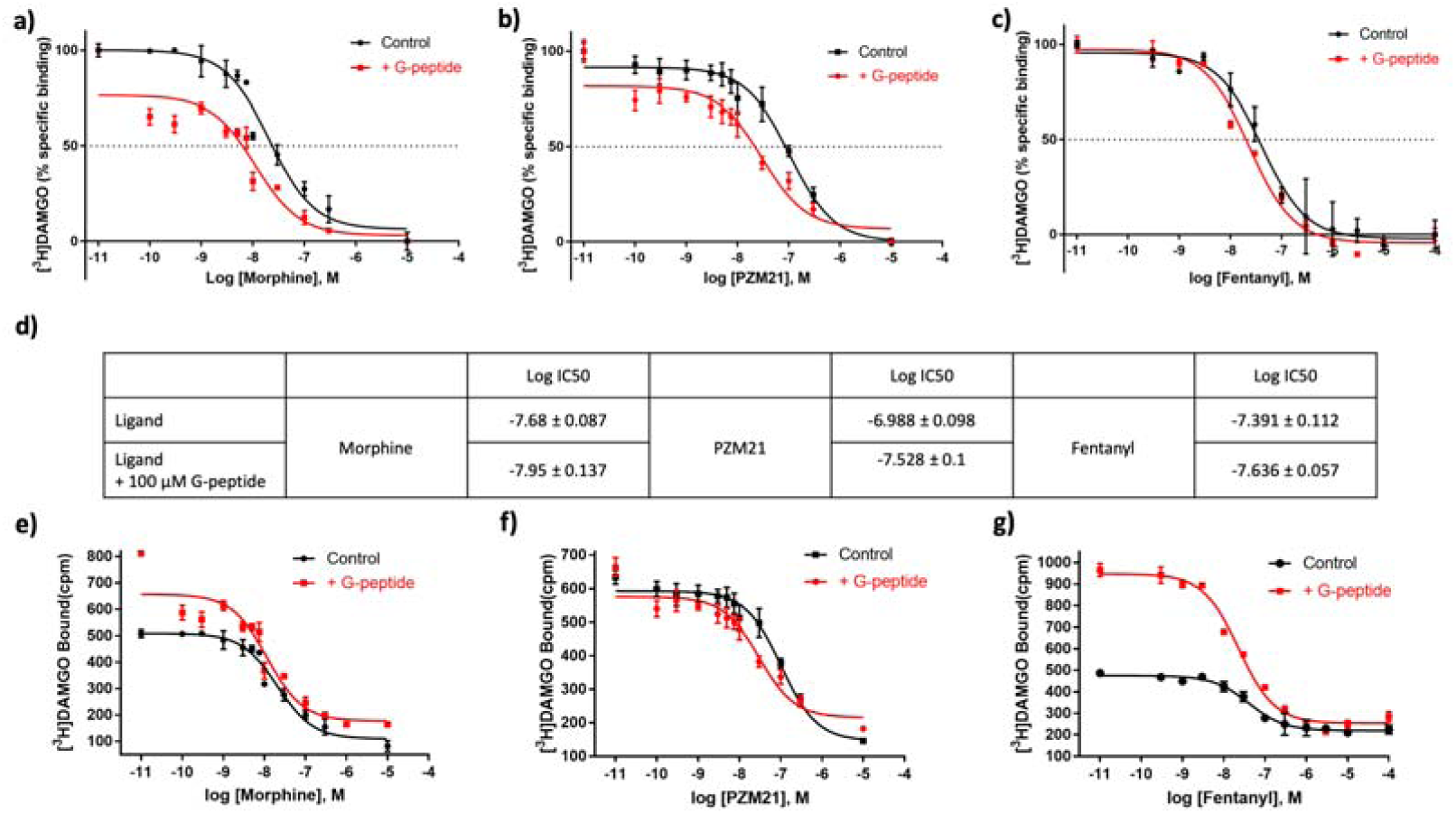
Detecting the interactions of the G-peptide with µOR. **a**, The effect of 100 µM G-peptide on the binding of morphine. The decrease in the specific binding percentage and leftward shift of the binding curve indicate the allosteric mode of the G-peptide interaction. **b**, The binding experiments performed for a G_i_-biased ligand, PZM21, show similar changes upon adding the peptide. **c**, Analogous experiments conducted for the β-arrestin-biased agonist fentanyl reveal a limited effect of the G-peptide on the binding characteristics of the ligand – a reduced shift of the binding curve and no effect on the specific binding of the radioligand [^3^H]DAMGO. **d**, The measured IC_50_ values for the experiments **a-c**. Data are representative of 5 independent experiments performed in duplicate. **e**, The binding experiments also revealed an increase in the total binding of [^3^H]DAMGO, correlating with previously published data. **f**, For PZM21, the total binding of the radioligand was not affected, but the baseline of the binding curve was elevated. **g**, The total binding of [^3^H]DAMGO is drastically increased in the experiments for fentanyl. This result in combination with limited binding affinity and nonspecific binding shifts suggests the intracellular binding site of the G-peptide, as it modulates the binding of the radioligand, not fentanyl.

### Computational study

In complement to the *in vitro* experiments, we studied the interactions between the G-peptide and µOR *in silico* to gain insight into the formation of the ternary complex. µOR has been crystallized in both active (PDB^22^ code: 5C1M^20^) and inactive (PDB code: 4DKL^23^) conformations and, recently, in complex with G_i_ (PDB code: 6DDE^24^). Unlike previous computational studies on GPCR-G protein complexes,^25,26^ where the starting point of the simulations was inferred from the crystal structure of the β_2_-AR-G_s_ complex,^27^ we performed simulations of the systems containing the receptor with docked morphine along with one G-peptide in close proximity to the intracellular binding site. The purpose of the simulations was to evaluate whether the peptide is likely to enter the crevice of µOR and whether it forms stable complexes with the receptor. We also performed simulations of the G-peptide in water, showing the oscillations between the helical and unstructured states (Supplementary Figure 4). For simulations with an inactive-state receptor, we observed the peptide approaching the binding site and making a network of contacts with µOR, with a number of hydrogen bonds between R^3×50^ and G.H5.22 (denoted in common numbering schemes for GPCRs^28^ and G proteins,^16^ respectively) as the main contributors (Figure 3c). A comparison of the conformation of the complex and recently released µOR-G_i_ complex revealed a similar angle of H5; however, the simulated G-peptide was translated by one helix turn due to the closed intracellular binding cleft (Figure 3a, b). For the same reason, the network of contacts, both van der Waals and hydrogen bonds, differed between the calculated and crystalized complexes (Figure 3c), suggesting two different conformational states of the receptor. This finding corresponds to the published model of high- and low-affinity ternary complexes with the G protein, where for the low-affinity complex, GTP bound to the Gα subunit, the binding of G.H5 is shallow, and upon release of GDP, the stable, closed active conformation of the GPCR is formed. We also observed a bend at the end of TM6 in the simulations, indicating the beginning of the conformational change in the receptor; however, the timeframe of the MD simulations (200 ns for production runs) did not allow us to observe the full transition into the active state. Interestingly, in the simulations with the active conformation of the receptor, we did not observe the G-peptide binding the receptor despite similar starting positions of the G-peptide (Supplementary Figures 5 and 6). The G-peptide would not enter the binding pocket even if pushed further into the binding cleft, resulting in interactions with ICL3 in the best-case scenario (Figure 3d, 3e; see an example in Supplementary Figure 6).

**Figure 3.**
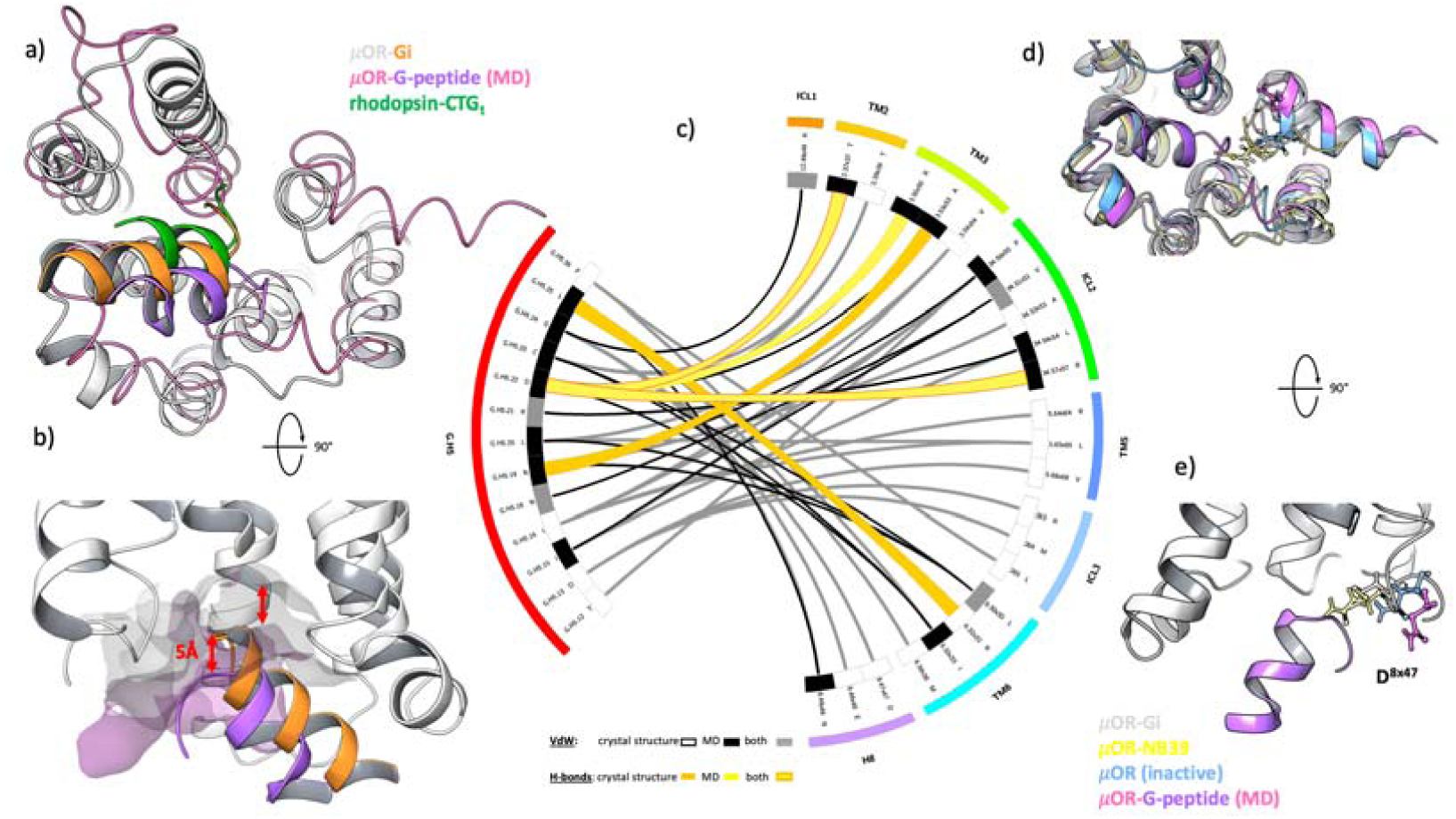
Analysis of the modelled µOR-G-peptide complex. **a**, The representative µOR-G-peptide complex obtained through MD simulations (pink ribbon, purple cartoon) overlapped with the crystal structures of metarhodopsin II crystalized with the C-terminal peptide of transducin (green cartoon, rhodopsin structure not shown for clarity, PDB code: 3PQR) and the µOR-G_i_ complex (white ribbons for the receptor and orange for C-terminal part of G_i_, PDB code: 6DDE), showing a similar angle of the complexed G-peptide. **b**, The difference in depth of binding between G_i_ and the G-peptide corresponds to the difference in the depth of intracellular binding sites of the active and inactive conformations of µOR. **c**, Comparison of the interaction network between G_i_ and µOR and the G-peptide and µOR. The bands in the inner circle indicate the contacts that are unique to the crystal structure (white), unique to the calculated complex (grey), or common (black). The main difference in specific interactions detected is the presence of hydrogen bonds between D^G.H5.22^ and R^3×50^, as found in the MD simulations (yellow ribbons), that are not present in the crystal structure (orange ribbons, shared hydrogen bonds are depicted with yellow ribbons with orange stroke). The change in the van der Waals network of contacts (crystal structure – grey lines, MD – black lines) is a consequence of the shallow binding pocket of the inactive-state µOR used in the experiments, resulting in a reduced solvent-accessible surface area of the crevice. **d**,**e**, Comparison of side-chain conformations of D^8×47^ in active-(µOR-NB39, PDB code: 5C1M – yellow ribbons and atoms; µOR-G_i_ as in **a**, side-chain of D^8×47^, missing in the electron density, has been modelled) and inactive-state (blue ribbons and atoms, PDB code: 4DKL; modelled complex as in **a**) conformations of µOR. In the inactive structures, the residue is facing away from the binding site, allowing the G-peptide to enter the pocket, whereas active structures have D^8×47^ facing towards the TM bundle, interacting with the G protein or a nanobody and stabilizing the complex but on the other hand preventing the G-peptide from entering the binding site in the simulations performed on the active receptor conformation.

## Discussion

### *In vitro* results

Our experiments show that the allosteric effect from a G protein to the agonist binding pocket, postulated by DeVree and Sunahara,^8^ can be mimicked by a synthetic peptide derived from the C-terminus of Gαi and detected in relatively simple competition binding experiments.

The saturation experiments performed with a radioligand and varying concentrations of the G-peptide showed the peptide affecting the binding in concentrations as low as 10 µM (Supplementary Figure 1); however, the experiments with morphine indicate that the characteristic shift in the binding curve associated with allosteric modulation cannot be observed when G-peptide concentrations are below 100 µM. This concentration corresponds to values reported for the rhodopsin study by Hamm *et al*. (100 – 700 µM range) but is significantly higher than the concentrations of nanobodies reported in the literature (5 µM). The reason for this discrepancy can be the promiscuity of the peptide – sharing the sequence with Gα_i_, it can potentially cross interact with approximately 200 other GPCRs coupled to G_i_, thus decreasing the bioavailability of the peptide.

In our opinion, µOR was the key element of the experiments. The variety of well-known ligands, the availability of functionally selective tool compounds and a number of crystal structures deposited in the PDB enabled the conduct and analysis of the experiments, both *in vitro* and *in silico*. In fact, it would be hardly possible to find another suitable target for such proof of concept.

### Computational study

Our approach to the computational study differs significantly from those published previously.^29–31^ Here, at the starting point of the MD simulations, the G-peptide was placed outside of the intracellular binding cleft in close proximity to the receptor (Supplementary Figure 5). This approach allowed us to observe unbiased binding of the peptide to µOR, as no peptide-receptor interactions were defined in the starting point of the simulation. Such a simulation system resulted in G-peptide drifting away from the receptor and failing to bind to the agonist-µOR complex. We have, however, introduced a bias on the secondary structure of the G-peptide, defining it on the basis of the C-terminus of the Gα_s_ co-crystallized with β_2_-AR.^19^ The native C-terminus of G proteins remains unstructured when not bound to a GPCR^32^, and our simulations on the G-peptide in water showed the instability of its secondary structure, oscillating between helical and unstructured (Supplementary Figure 4). Using the unstructured G-peptide as a starting point would significantly increase the simulation time, and the aim of the MD simulations run here was to detect the binding event of the G-peptide to the µOR. On the other hand, the helical structure of the peptide remained stable throughout the simulations of the ternary complex.

Binding of the G-peptide to the receptor, as revealed by the MD simulations, starts with forming a hydrogen bond between R^3×50^ and D^G.H5.22^, followed by the formation of a hydrogen bond with ICL2 (R^34×57^), stabilizing the orientation of the G-peptide within the binding cleft. This observation follows the findings of Elgeti *et al*.,^30^ which proposed a similar binding mechanism for rhodopsin and transducin CT-peptide. There, R^3×50^ served as the anchor point for the peptide, leading, through mutual conformational adjustments, to stabilization of the helical structures of both the peptide and ICL3 and thus formation of the stable active conformation of rhodopsin. Those conclusions were drawn from NMR studies accompanied by MD simulations of crystal structures of rhodopsin (inactive, active and active with G_t_ peptide), while in our experiments, we observed the same sequence of events starting from receptor and G-peptide separation.

The final G-peptide conformation obtained from MD simulations differs from the conformation of the corresponding region of Gα_i_ from the crystal structure (PDB code: 6DDE) as well as from the conformation of transducin peptide crystallized with rhodopsin (PDB code: 3PQR). The angle at which the G-peptide remains in the binding site is similar to that in both crystal structures; however, it is rotated by approximately 90°. In addition, the G-peptide does not enter as deep into the intracellular side of the receptor as can be observed in the crystal structures. The reason for this result is the depth of the binding site of the inactive-state µOR used in the calculations. Through the course of the MD simulations, we were not able to recreate the full transition to the active conformation of µOR, as the time scale of that rearrangement is significantly larger than that of the simulations performed. The different depth of binding of the G-peptide indicates differences in the interaction network between peptide and receptor (Figure 3). Hydrogen bonds with TM6 (R^6×32^ – L^G.H5.25^) are missing from the simulation results; TM3 also interacts through A^3×53^ – N^G.H5.19^, and the hydrogen bond between R^3×50^ and D^G.H5.22^ is absent in the crystal structures. The presence of key anchor hydrogen bonds with R^3×50^ but a lack of hallmarks of GPCR activation (rotation of TM6 as well as wide opening of the intracellular crevice) indicate that the G-peptide-receptor conformation we obtained from MD simulations corresponds to the “low-affinity state” described by DeVree *et al*. or the early engagement complex described by Elgeti *et al*.

An interesting observation comes from the simulations performed with the active-state µOR (PDB code: 5C1M) as a starting point. In all of the MD runs we performed, the G-peptide failed to enter the binding site of the receptor and was repelled by the interaction with D^8×47^ from the hinge region between TM7 and H8 of the receptor. This result indicates that the G-peptide-receptor early engagement complex, or “low-affinity state”, is essential for transition into the fully active state of the GPCR. The G protein (or G-peptide in this case) does not find an active-state receptor that it can stabilize. It engages the agonist-bound GPCR and pushes it into an activated “high-affinity state”, which it then stabilizes. These findings provide evidence for the reaction mechanism driving GPCR activation described by Elgeti *et al*. but also add constraints to it, showing that formation of the early engagement complex with a characteristic hydrogen bond between the G-peptide and R3×50 for µOR is the first and essential step in the process of receptor activation.

## Conclusions

Our results demonstrate that experiments with rhodopsin and the C-terminal peptide of a G protein performed nearly 30 years ago can be extrapolated to µOR. Moreover, our hypothesis of the length of ICL3 being the caveat of this approach can explain the lack of literature reports for the targets of interest of the GPCR research community. There is an overrepresentation of receptors with an extended ICL3 among GPCRs, but our findings open nearly 200 other receptors for further investigation of activation and biased signalling using synthetic peptides. Indeed, we show that by using the G-peptide, it is possible to differentiate between functionally selective compounds without a need for either cell-based assays detecting secondary messengers or receptor-tailored antibodies.^33^ The computational study performed here corresponds to the model of rhodopsin activation, proposing a cooperative process of the receptor conformational rearrangement, indicating that it can indeed be universal for the GPCR superfamily. This study also conforms to the low- and high-affinity state model, showing that the initial low-affinity G-peptide-receptor ensemble is essential for the transition into a high-affinity ternary complex.

## Acknowledgements

We would like to thank X. Deupi, A. S. Hauser, K. P. Hofmann, A. J. Kooistra and F. S. Jørgensen for valuable comments, insights and discussion.

The study was supported by the grant PRELUDIUM 2015/19/N/NZ2/01818 financed by the National Science Centre, Poland.

## Methods

### Sequence analysis

The G protein binding data were extracted from GPCRdb.^1^ The GPCRdb-annotated sequence segments were also extracted with an in-house script.

### Molecular dynamics simulations

The structure of mouse µOR pre-aligned to a membrane was retrieved from the OPM database (entries 4DKL for inactive and 5C1M for active receptor conformations). The structure was then prepared with Schrödinger’s Protein Preparation Wizard (bond orders, charges and OPLS3 force field parameters). Missing side chains as well as missing residues of ICL3 (residue numbers 264-269) were rebuilt with Prime.^2^ The morphine structure was prepared using LigPrep^3^ and docked into the prepared receptor (Schrödinger Glide XP, default settings).^4^

A homology model of the G-peptide was prepared to retain the helical conformation. The peptide of 12 residues was built on the template 3SN6 (chain A) and prepared with Protein Preparation Wizard (charges and OPLS3 force field parameters as well as terminal acyl and amide groups). The crystal structure of the β_2_-adrenergic receptor bound to G_s_ was the only available structure with a fully folded G.H5 of a G protein. The best crystal structure of Gα_i_, 1BOF, has the C-terminus partially unfolded and thus is unsuitable for the experiments.

Two variants of the peptide were tested: wild type with 354F (G.H5.26) as the C-terminus and a reversed version, with 354F as the N-terminus. We did not observe any interactions between cap-protecting groups and the receptor in the MD simulations, and both variants of the peptide behaved in a similar manner.

Models were created in Schrödinger’s Maestro, using Prime backend (homology modelling) along with Protein Preparation Wizard (acylation and amidation).

The simulation system consisted of the receptor, the peptide and a POPC membrane. Cl^-^ ions were added to neutralize the system. The total atom count of the simulation system was approximately 47 000.

The G-peptide was manually placed in close proximity to the intracellular binding cleft of the receptor. We ran several test simulations with the peptide placed further from the receptor to observe if spontaneous binding would occur. We observed spontaneous binding; however, this approach was not optimal for investigation of the µOR-G-peptide complex, and for these runs, the starting point for the simulations was with the two molecules close to each other. For these runs, which were repeated five times, we could observe the binding of the G-peptide in 4 out of 5 experiments with an inactive conformation of µOR. The starting conformation is visualized in Extended Data Figure 3.

Relaxation of the MD systems followed Schrödinger’s relaxation protocol for membrane proteins. Production runs were performed at 310 K using the NPAT ensemble. The simulation time was 200 ns per simulation. Schrödinger software and in-house scripts utilizing Schrödinger python libraries were used for analysis of the trajectories. The last frame of a representative MD run was used to construct the interactions plot in Figure 3c. The plot was generated with Circos.^5^ Structure figures were prepared with Schrödinger’s Maestro 2018-1.

### Peptide synthesis

All reagents, amino acids and solvents were purchased from commercial suppliers and used without further purification.

The G-peptide was synthesized manually by a standard method, 9-fluorenyl-methoxycarbonyl (Fmoc)-based SPPS on Rink amide resin in DMF, using an HBTU/HOBt/DIPEA mixture for coupling and piperidine for deprotection steps. The G-peptide was cleaved from the peptidyl resin by TFA, purified by RP-HPLC and verified by ESI-MS.

The G-peptide was characterized with a Shimadzu instrument: liquid chromatograph connected with a mass detector (LC/MS) using a Jupiter 4 µm Proteo 90 Å C12 column (250 × 4.6 mm, 4 µm; Phenomenex, USA) with a flow rate for LC of 1.2 ml/min and for MS of 0.4 ml/min. Analysis was performed using a linear gradient of 10 to 30% B over 10 min (A: 0.05% FA in H_2_O, B: 0.05% FA in ACN). Detection was performed at 210 nm. The G-peptide was purified on a preparative Shimadzu HPLC system with a reversed-phase Jupiter® 10 µm Proteo 90 Å (250 × 21.2 mm), AXIA™, C12 column at a flow rate of 20 ml/min using a linear gradient of 20 to 35% of B over 35 min (A: 0.1% TFA in H_2_O, B: 0.1% TFA in ACN).

### Radioligand-binding assay

Crude membrane preparations isolated from Wistar rat brains were incubated at 25°C for 60 min with 0.5 nM [^3^H]DAMGO in a total volume of 1 ml of 50 mM Tris–HCl (pH 7.4) containing bovine serum albumin (BSA) (1 mg/ml), bacitracin (50 mg/ml), bestatin (30 mM) and captopril (10 mM). All reactions were carried out in duplicate at a 10 mM peptide concentration. Incubations were terminated by rapid filtration through GF/B Whatman glass fibre strips using a Brandel 24 Sample Semi-Auto Harvester. The filters were washed with 2 ml of an ice-cold saline solution, and the bound radioactivity was measured in a MicroBeta LS, TriLux liquid scintillation counter (PerkinElmer). Nonspecific binding was determined in the presence of naltrexone hydrochloride (10 mM). The data were analysed by a nonlinear least square regression analysis computer program GraphPad Prism.

We have investigated the effect of the G-peptide on [^3^H]DAMGO binding to provide evidence on the allosteric cooperativity between the two. We conducted two types of experiments: competition binding against [^3^H]DAMGO and saturation binding of the radioligand. The competition experiment allowed for determination of the IC_50_ of the G-peptide for the high and low range of concentrations, with values of 27.3 and 11.55 µM, respectively (see Supplementary Figure 1 for details). These values, compared to the nanomolar activity of DAMGO, render the G-peptide inactive through the orthosteric binding site of µOR. On the other hand, the saturation experiment revealed a concentration-dependent decrease in the B_max_ value of DAMGO upon adding different concentrations of the G-peptide (two sets of experiments with G-peptide concentration ranging from 0.01–25 µM and 0.01– 200 µM, see Supplementary Figure 2). These results indicate the allosteric coupling between DAMGO and the G-peptide and strengthen the interpretation of the binding curve shifts presented in Figure 2 as the allosteric effects.

