## Supplementary Figures for "Gα_i_-derived peptide binds the µ-opioid receptor"

### Supplementary Figure 1

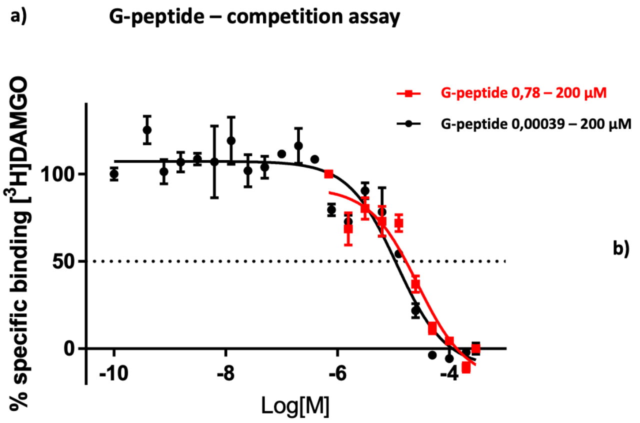


Results

| | | |
|---|---|---|
| Log IC50 | -4,564 | -4,938 |
| IC50 [µM] | 27,3 | 11,55 |

b)

| Concentrations [µM] | |
|---|---|
| 200 | 0,39 |
| 100 | 0,2 |
| 50 | 0,1 |
| 25 | 0,05 |
| 12,5 | 0,025 |
| 6,25 | 0,00625 |
| 3,125 | 0,003125 |
| 1,5625 | 0,001563 |
| 0,78125 | 0,000781 |
| | 0,000391 |

**Supplementary Figure 1 | Evaluation of the orthosteric effect of the G-peptide a,b,** The results of the competition binding assay for the G-peptide show no significant orthosteric coupling to µOR. The $IC_{50}$ values at the micromolar level (**a**) determined at two concentration ranges (**b**) are vastly higher than the subnanomolar activity of the reference ligand DAMGO (Handa 1981). These experiments contribute to the hypothesis that the effects observed for subsequent addition of the G-peptide are of allosteric nature.

### Supplementary Figure 2

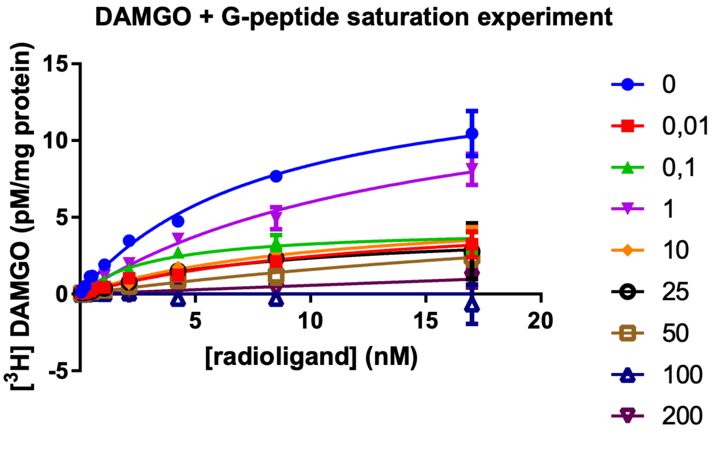


| G-peptide concentration [µM] | 0,01 | 0.1 | 1 | 10 | 25 | 50 | 100 | 200 |
|---|---|---|---|---|---|---|---|---|
| $B_{max}$ | 5.627 | 4.313 | 14.91 | 5.711 | 4.317 | 8.021 | n.d. | n.d. |
| $K_d$ | 13.02 | 3.236 | 14.86 | 10.77 | 8.424 | 40.25 | n.d. | n.d. |

**Supplementary Figure 2 | Saturation experiments for DAMGO and the G-peptide.** The addition of varying concentrations of the G-peptide to the experiment resulted in a change in both the equilibrium dissociation constant $K_d$ and $B_{max}$. For the high concentrations of G-peptide (namely, 100 and 200 µM), we were not able to reliably fit the binding model.

### Supplementary Figure 3

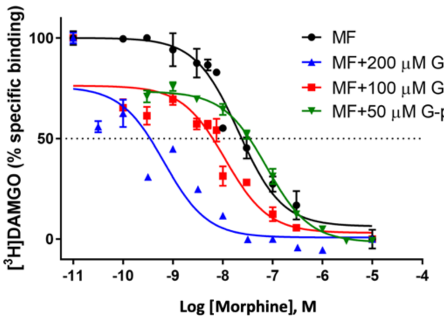


| | Log IC50 | IC50 (nM) |
|---|---|---|
| Morphine | -7.68 ± 0.087 | 21 |
| Morphine + 50 μM G-peptide | -7.10 ± 0.044 | 77 |
| Morphine + 100 μM G-peptide | -7.95 ± 0.137 | 11 |
| Morphine + 200 μM G-peptide | -9.2 ± 0.2 | 0.66 |

**Supplementary Figure 3 | Binding experiments for morphine with three different concentrations of the G-peptide**. The addition of various concentrations of the G-peptide to the binding experiment for morphine resulted in a dose-dependent shift of the binding curve. The addition of 200 μM G-peptide resulted in a 30-fold increase in morphine affinity, corresponding to the values reported for experiments with nanobodies. The results, however, yield high uncertainty, possibly due to peptide aggregation caused by its high concentration. For this reason, further experiments were conducted with the peptide concentration of 100 μM, giving more consistent results even though the affinity shift was lower.

Supplementary Figure 4

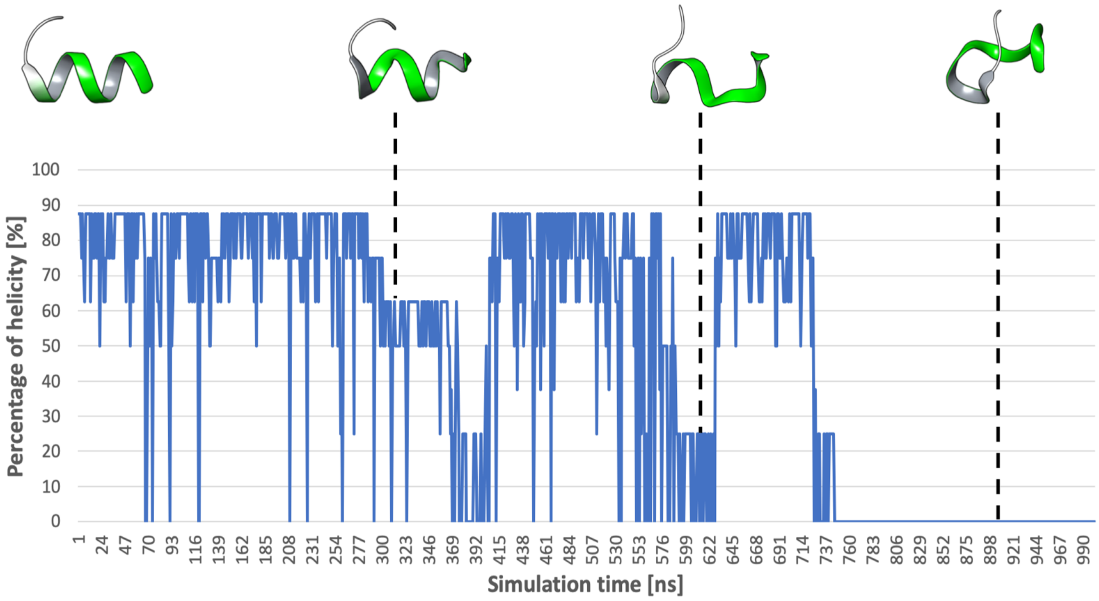


**Supplementary Figure 4 | MD Simulation of the G-peptide in water.** We performed a 1000 ns-long simulation of the G-peptide in water to evaluate the stability of its secondary structure. We measured the helicity of ten residues (highlighted in **green**) across the simulation. The results show that the peptide oscillates between helical and unstructured conformations. These results are in line with literature data, indicating that the C-terminus of the G protein is intrinsically disordered when outside of the intracellular binding cleft of the GPCRs.

Supplementary Figure 5

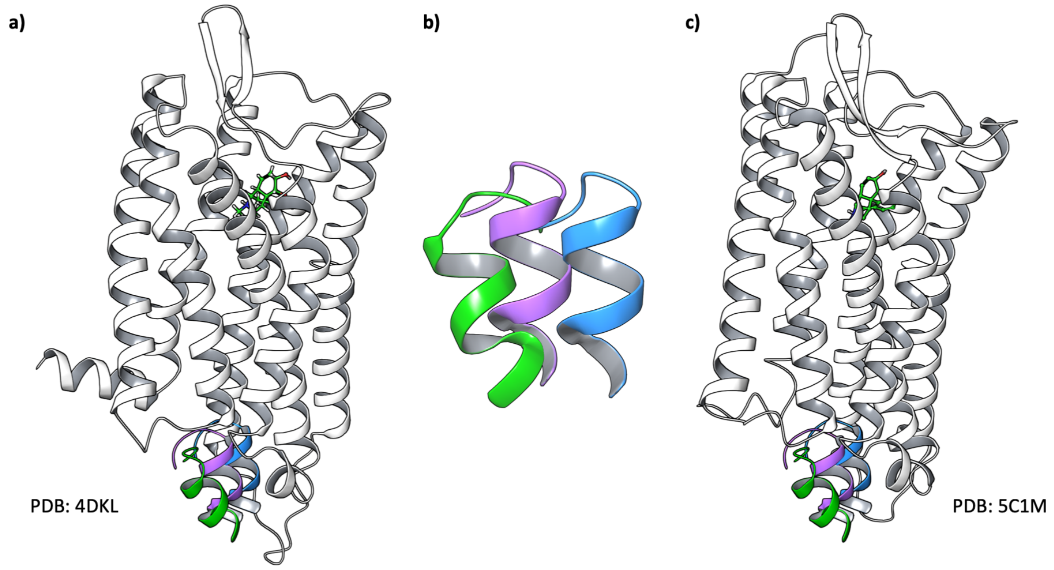


**Supplementary Figure 5 | Starting points for the MD simulation experiments.** The model of the G-peptide was manually placed in the vicinity of the G protein binding site of the receptor in a way that no contacts were between the peptide and the receptor at the start of the simulation. **a,** The starting systems for the inactive conformation of μOR (PDB code: 4DKL) with a molecule of morphine docked into the orthosteric site. **b,** The different orientations of the G-peptide used in MD simulations. Not only was it translated but different rotations were also considered. **c,** The system setup for the active conformation of μOR (PDB code: 5C1M) with a molecule of morphine docked into the orthosteric site. In both setups, **a** and **c**, the receptor sequence was reverted to that of the human wild type. Missing residues, including ICL3 of the inactive structure, were rebuilt into the model.

### Supplementary Figure 6

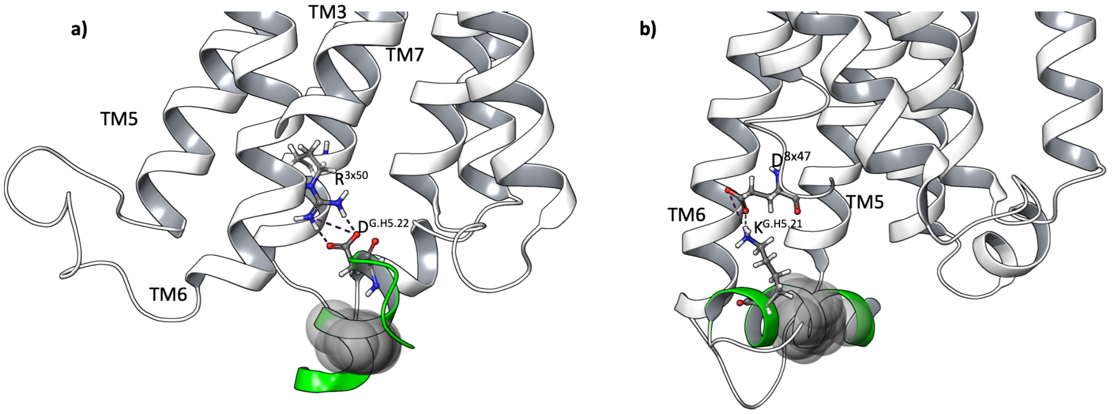


**Supplementary Figure 6 | The final conformations of the receptor-G-peptide complexes.** The grey spheres are the trace of the centre of mass of the G-peptide throughout the MD simulation (20 spheres sampled from the trajectory are displayed). **a,** The results of the MD simulation with the inactive conformation of μOR (PDB code: 4DKL). The G-peptide (green ribbon) forms a stable hydrogen bond with $D^{3x50}$ (D**R**Y motif) that stabilizes it within the binding site and limits the movement of the peptide (grey spheres). **b,** The simulation for the active-state conformation of μOR (PDB code: 5C1M) does lead to different results. The movement of the G-peptide is less restrained because it is unable to enter the intracellular binding site due to a hydrogen bond with $D^{8x47}$. Part of TM7 and H8 were removed for clarity in both **a** and **b**.
